# Quantitative Delineation of Herpesviruses in Bats for use in Ecological Studies

**DOI:** 10.1101/856518

**Authors:** Anna R. Sjodin, Michael R. Willig, Simon J. Anthony

## Abstract

Public health concerns about recent viral epidemics have motivated researchers to seek transdisciplinary understanding of infection in wildlife hosts. With its deep history devoted to explaining the abundance and distribution of organisms, ecology can augment current methods for studying viral dynamics. However, datasets allowing ecological explorations of viral communities are lacking, and common methods for delineating viral operational taxonomic units (OTUs), or “species”, are subjective. Here, we comprehensively sampled 1,086 bats from two Puerto Rican caves and tested them for infection with herpesviruses. Using percent identity of nucleotides and a machine learning algorithm, we categorized herpesviruses into 41 OTUs, representing approximately 80% of all herpesviruses in the host community. Although 13 OTUs were detected in multiple host species, OTUs generally exhibited host specificity by infecting a core host species at a significantly higher prevalence than in all other species combined. Only two OTUs showed significantly different prevalence between host sexes. This work is the first exploration of viral community ecology in a community of wildlife hosts.

## Introduction

Public health concerns associated with recent and ongoing outbreaks of viruses (e.g., Ebola virus and Zika virus) have motivated researchers to investigate viral diversity in wildlife hosts from previously unexplored perspectives. Ecology has a long history of quantifying the abundance and distribution of organisms and has been increasingly integrated into viral studies (Anthony et al. 2015, Seabloom et al. 2015). Traditionally, such efforts have focused on isolated cases of single host-single pathogen relationships (Telfer et al. 2010). However, infected host individuals generally have multiple pathogens, and infection with one pathogen can affect infection, virulence, or disease emergence of the others (Pedersen and Fenton 2006, Jolles et al. 2008, Telfer et al. 2010, Seabloom et al. 2015). This suggests that perspectives and approaches from community ecology can complement current methods in virology.

Three major challenges exist when integrating community ecology and virology. First, most viruses are rare (i.e. exist at low prevalence; Tang et al. 2006, Winker et al. 2008, Wacharapluesadee et al. 2010, Anthony et al. 2015), so the probability of infection in a host individual is low. Consequently, large datasets of comprehensively sampled host populations are needed to address high levels of uncertainty associated with low viral prevalence. Few such datasets exist. Second, most wildlife viruses are unknown to science (Anthony et al. 2013, Anthony et al. 2017, Carlson et al. 2019), and the process of defining viral species is time-consuming (King et al 2012). This means that waiting for official designations of viral species for use in community-level analyses does not address the urgency of current health challenges posed by wildlife viruses. Alternatives to official designation of viral species include delineating operational taxonomic units (OTUs) using monophyletic groups (Anthony et al 2015) or percent identity histograms (Maes et al. 2009, Anthony et al. 2017). However, both such methods rely on subjectively chosen cutoff points to differentiate OTUs. Finally, many viral infections last only hours to days, making detection difficult (Klenk et al. 2004, Lee et al. 2009, Henaux and Samuel 2011, King et al. 2012, Vetter et al. 2016).

Herpesviruses are large, enveloped, double-stranded DNA viruses that infect all vertebrate classes and mollusks (King et al. 2012, Azab et al. 2018). In contrast to many viruses, herpesviruses generally occur at high prevalence (Cone et al. 1993, Kidd et al. 1996, Cortez et al. 2008, Imbronito et al. 2008, Tenorio de Franca et al. 2012, Phalen et al. 2017, Tazikeh et al. 2019), and they establish latent, long-term (often for the entirety of the host’s life) infections (King et al. 2012). Using herpesviruses as the subject of ecological analyses can therefore minimize two challenges that stymie advancements in viral ecology.

Herpesviruses were first detected in bats in 1996 (Tandler 1996). Subsequent work shows that bats likely played a critical role in the evolution and diversification of herpesviruses (Escalera-Zamudio et al. 2016). Bats generally have a negative reputation as carriers of deadly viruses that cause human diseases such as rabies, Nipah, and SARS (Wibbelt et al. 2010), and consequently, most research on bat viruses has focused on these pathogens of concern for public health. Although a number of studies have also detected herpesviruses in bats (Wibbelt et al. 2007, Razafindratsimandresy et al. 2009, Zhang et al. 2012, Sasaki et al. 2014, Pozo et al. 2016, Zheng et al. 2016, Holz et al. 2018, Wada et al. 2018), little is known about viral shedding, host specificity, or the diversity of herpesviruses in bats (Zheng et al. 2016).

Herpesviruses are generally considered to be host-specific (King et al. 2012). However, inter-host transmission has been documented when close contact occurs between individuals of different species, especially when host species are closely related (e.g. within primates: Weigler 1992; within turtles: Greenblatt et al. 2005; within ungulates: Russell et al. 2009). Multiple bat species come into close proximity within caves where they roost, presenting an opportunity for cross-species viral dispersal. Indeed, cross-species transmission and host switching are common, and have likely been the norm throughout the evolutionary history of the herpesvirus lineage, especially in bat hosts (Escalera-Zamudio et al. 2016, Zheng et al. 2016, Azab et al. 2018, Wada et al. 2018).

The first goal of this research was to intensively sample bat communities in two Puerto Rican caves, with the intent of saturating viral discovery of herpesviruses for use in ecological analyses. The second goal was to determine a quantitative technique to differentiate operational taxonomic units (OTUs) of herpesviruses as a fast, objective alternative to defining viral species. Finally, we aimed to describe patterns of host preference for herpesviruses. Given recent advancements in the understanding of host sharing throughout the history of the herpesvirus lineage, we hypothesized that particular OTUs would occur in multiple bat species but would have significantly higher prevalence in one of the species (i.e. viruses will have a core host species; Bush and Holmes 1986); in all satellite (i.e. “secondary”) hosts combined, viruses will have lower prevalence.

## Materials and Methods

### Sample Collection

Field work was conducted at two adjacent caves on the Mata de Plátano Nature Reserve in Arecibo, Puerto Rico (18° 24.868’ N, 66° 43.531’ W). The 13 bat species on the island are well-documented and have been studied extensively (Gannon et al. 2005). Mata de Plátano Nature Reserve (operated by InterAmerican University, Bayamon, Puerto Rico) is in the north-central, karstic region of the island, an area dominated by a multitude of caves that are suitable for roosting bats.

Culebrones Cave is structurally complex and hot, with temperatures reaching 40° C and relative humidity at 100%. It is home to an estimated 300,000 individual bats representing six species: *Pteronotus quadridens, P. portoricensis, Mormoops blainvillii, Monophyllus redmani, Erophylla sezekorni*, and *Brachyphylla cavernarum* (Rodríguez-Duran 2009). Bats were sampled for 28 nights between June and August 2017. A harp trap was placed at sunset (approximately 18:00) immediately outside of the opening, and monitored continually.

Larva Cave is cooler (ambient temperature), smaller, and less structurally complex compared to Culebrones Cave. It hosts a small total number of bats (30-200; ARS personal observation) representing two species, *Artibeus jamaicensis* and *Eptesicus fuscus*. Bats were sampled from Larva Cave on seven instances between June and August 2017, using two different techniques. After sunset, mist nets were placed along a trail outside of the cave entrance to catch exiting bats and were checked at least every ten minutes. Hand nets were also used to capture bats roosting inside the cave.

Regardless of sampling method, each captured bat was placed into a porous cotton holding bag. Individuals were identified to species following Gannon et al. (2005), and a clean cotton-tipped swab was used to collect saliva from the bat’s mouth. Swabs were placed in individual cryovials containing viral transport medium and sent to Columbia University’s Center for Infection and Immunity, in a dry shipper, and stored at −80°C. All methods were approved by the University of Connecticut Institutional Animal Care and Use Committee (IACUC, protocol A15-032).

### Viral Screening

Total nucleic acids were extracted from each swab using the EasyMag platform (bioMerieux, Inc.), and cDNA was synthesized using SuperScript III first-strand synthesis supermix (Invitrogen). Nested consensus polymerase chain reaction (cPCR) was performed twice on each sample, targeting a region of the highly conserved catalytic subunit of the DNA polymerase gene (Van deVanter et al. 1996). cPCR is a broadly reactive method that allows detection of novel viruses by using nonspecific primers targeted at viral families or genera (Anthony et al. 2013, 2015). A constructed synthetic plasmid was used as a positive control to confirm effective implementation of the assay and to detect possible contamination. PCR gel products (1% agarose) of expected size were cloned into Strataclone PCR cloning vector, and twelve colonies were sequenced to confirm identity and detection of co-occurring viruses. Each clone was cross-referenced using the GenBank nucleotide database to confirm that sequences were indeed herpesviruses. Sequence data will be available at github.com/ARSjodin upon publication.

### OTU Delineation

A nucleotide alignment of 3,131 herpesvirus clones was conducted using the ClustalW alignment in Geneious version 11.0.5, and a pairwise percent identity (PID) matrix was calculated based on percent similarity of nucleotides in each sequence pair. A PID histogram was then generated (Maes et al. 2009). To evaluate the robustness of cutoffs to errors in alignment, the R package Biostrings (Pages et al. 2017) was used to conduct pairwise alignments of clones. A second PID histogram was generated, and frequency clusters were compared between the two PIDs. The number of OTUs was estimated using the cutoff extremes for each level of classification, based on total alignment and pairwise alignment (74, 81, 90 and 93 PID cutoffs, see Results and Figure 1).

**Figure 1:**
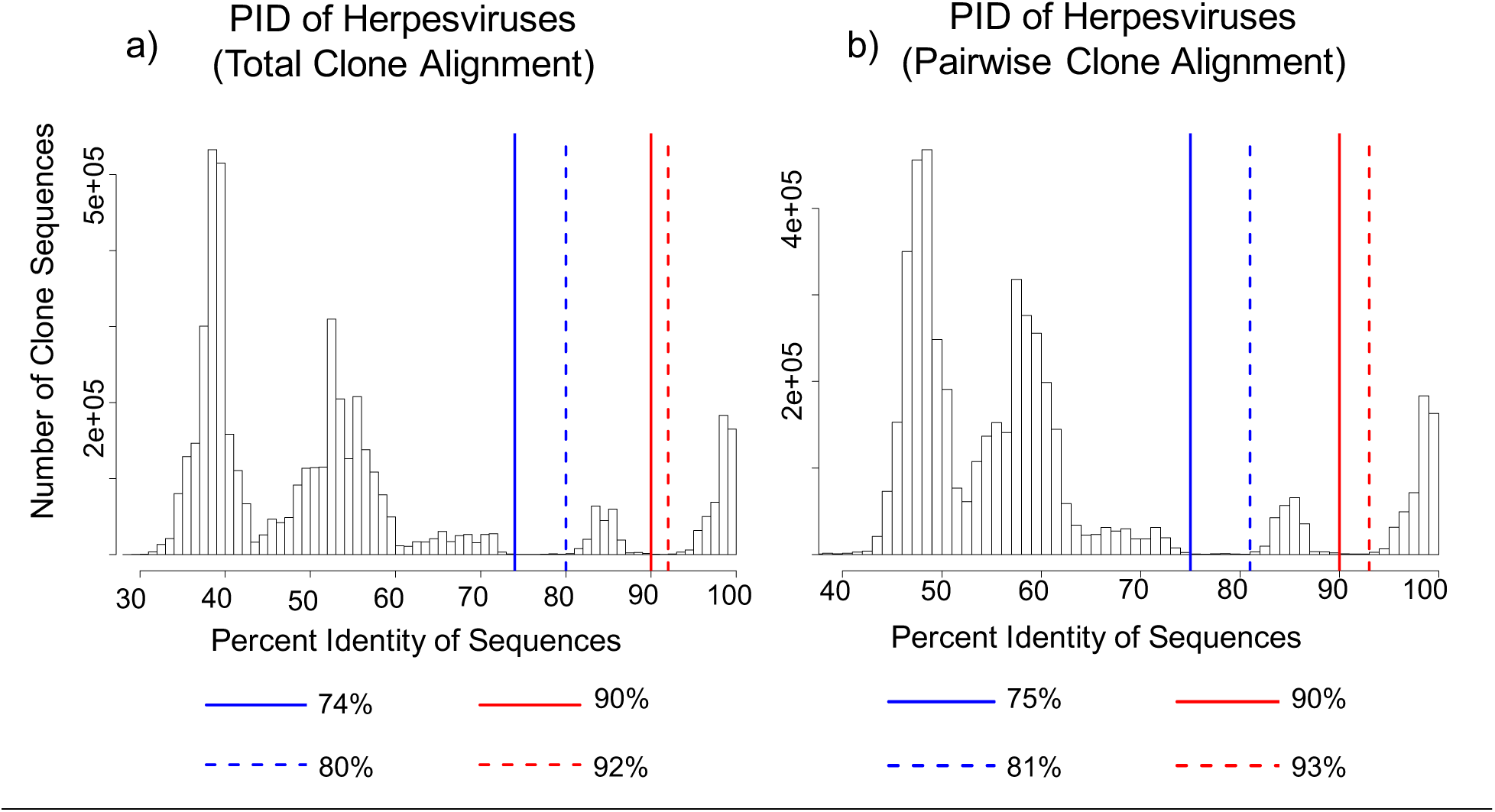
Percent Identity (PID) histograms of herpesviruses using the alignment of all clones (a) and pairwise alignment of clones (b). The x-axis represents the percent identity of clone sequences compared pairwise. The y-axis represents the number of clone sequences. Vertical lines represent cutoff points corresponding to two distinct peaks, with blue lines representing the higher-level cutoff and the red lines representing the lower-level cutoff. Solid lines represent the low end of the cutoff ranges, and the dashed lines represent the upper end of the cutoff ranges. The distinct peaks are consistent regardless of alignment method.

To statistically support which OTU cutoff to use in subsequent analyses, the R package apcluster (Bodenhofer et al. 2011) was used to perform affinity propagation on the PID matrix of the total clone alignment. Affinity propagation is an algorithm that examines all data points in a similarity matrix to identify exemplars (Frey and Dueck 2007). The algorithm then clusters remaining data around an optimum number of exemplars by iteratively exchanging two types of “messages” between data points. In the context of viral sequences, the first type of message accumulates evidence for sequence *k* as an exemplar for sequence *i*, based on genetic distance between *k* and *i* (Frey and Dueck 2007). The second type of message reflects how many sequences besides *i* support *k* as the best choice of exemplar (Frey and Dueck 2007). When inputting the PID matrix of all clones, the number of clusters identified through affinity propagation can be interpreted as an optimal number of viral OTUs. Although this method has been used to define clusters of a single viral species (Fischer et al. 2017), this is the first instance in which it has been used to delineate viral “species” for the purpose of community-level analyses. To conduct affinity propagation analyses, the apcluster() command was run using the R package apcluster (Bodenhofer et al. 2011), with default values of input parameters (input data = NA, input preference = NA, q = 0.5, convits = 100, maxits = 1000, lam = 0.9). Finally, OTUs were refined manually based on PID and affinity (see Results).

Finally, sequences representing the polymerase gene of formally described beta-and gammaherpesvirus species, as defined by the International Committee on the Taxonomy of Viruses (ICTV; Order Herpesvirales Figure 5, King et al. 2012) were retrieved from GenBank. Matrices comparing PID of defined viral species were calculated for comparison to empirical values. For ease of exposition, “species” will be used moving forward to reference host taxa or formally described viral taxa, whereas “OTU” will be used to reference empirical viral taxa.

### Viral Discovery Curves

To determine how well the collected samples represent the metacommunity of herpesviruses infecting bats at Mata de Plátano Nature Reserve, viral discovery curves were created using the iNEXT package in R (Hsieh et al. 2016). By defining the metacommunity as all of the herpesviruses infecting all of the host species at Mata de Plátano Nature Reserve, an assumption is made that herpesviruses are shared between caves and among host species (i.e. no host specificity). Because this may not be a valid assumption, separate curves were generated for each bat species, as well as for each cave. Additionally, Chao2 estimates (Chao 1987) were calculated for the viral metacommunities of all bats at Mata de Plátano Nature Reserve, as well as for each cave and for each bat species separately.

### Host Preference

For each OTU that infected more than one host species, the host specificity index S_TD_* was calculated (Poulin and Mouillot 2005) as

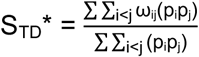

where *i* and *j* represent index values of host species, *p* represents OTU prevalence within host populations, and ω_ij_ is the phylogenetic distinctiveness of hosts. To measure phylogenetic distinctness, hosts of the same genus are assigned a value of one, hosts of the same family but different genera are assigned a value of two, and hosts of different families have a value of three (Poulin and Mouillot 2005). S_TD_* thus weighs host preference by the relatedness of hosts (incorporating evolution) and the prevalence of the pathogen in each of its hosts (incorporating ecology). The extent of viral sharing among hosts species was further examined by determining whether particular OTUs had a core host species. A core host species was defined as a single host species for which the OTU had significantly higher prevalence than in all other host species combined. To do this, the prevalence of each OTU in its core host and its combined prevalence in all other hosts were compared using Fisher’s exact test (Wassertheil-Smoller and Smoller 2015). For each OTU that infected more than one host individual, viral preference for host sex was examined by comparing OTU prevalence between male and females of the core host species, using Fisher’s exact test. All analyses were done using R version 3.4.3, and code used for all methods can be found at github.com/ARSjodin.

## Results

### Viral Screening

Oral swabs were collected from 1,086 bats representing eight species (*A. jamaicensis* and *E. fuscus* in Larva Cave, and *P. quadridens, P. portoricensis, Mor. blainvillii, Mon. redmani, E. sezekorni*, and *B. cavernarum* in Culebrones Cave; Table 1). Three hundred and thirty host individuals tested positive for herpesvirus, for a community-level prevalence of 30.4% (Table 1). Prevalence was similar in Culebrones (30.0%) and in Larva (35.3%) Caves. Herpesvirus prevalence at the host population-level ranged from 12.9% in *P. portoricensis* to 42.1% in *A. jamaicensis*. At least one individual from each host species tested positive, with the exception of *E. fuscus* (0/11). In females, prevalence ranged from 12.5% in *P. portoricensis* to 41.9% in *Mor. blainvillii*. In males, prevalence ranged from 14.3% in *P. portoricensis* to 42.4% in *A. jamaicensis*.

**Table 1:** Summary of capture numbers and virus screening results. Numbers of host individuals sampled (N), number testing positive for herpesvirus (N positive), and prevalence of herpesviruses are calculated for each host species and for each cave, as well as separately for males and females of each host species. Number of herpesvirus OTUs detected empirically (S) and estimate of herpesvirus OTU richness (Chao2), are compared to estimate the percentage of herpesvirus OTUs that was detected in this study (% completeness). No male *Eptesicus fuscus* were captured, so NA values are produced.

| Host Species | N (Males, Females) | N Positive (Males, Females) | Prevalence (Males, Females) | S (% Completeness) | N Common, N Rare | Chao2 (95% lower, upper) |
| --- | --- | --- | --- | --- | --- | --- |
| *Pteronotus quadridens* | 288 (155, 133) | 56 (39, 17) | 19.44 (25.16, 12.78) | 15 (55) | 5, 10 | 27.207 (17.194, 82.937) |
| *P. portoricensis* | 31 (7, 24) | 4 (1, 3) | 12.90 (14.29, 12.50) | 3 (93) | 2, 1 | 3.242 (3.013, 7.622) |
| *Mormoops blainvillii* | 299 (194, 105) | 120 (76, 44) | 40.13 (39.18, 41.90) | 9 (67) | 3, 6 | 13.485 (9.493, 49.788) |
| *Monophyllus redmani* | 262 (104, 158) | 90 (32, 58) | 34.35 (30.77, 36.71) | 11 (81) | 2, 9 | 13.656 (11.375, 29.801) |
| *Erophylla sezekorni* | 74 (42, 32) | 12 (8, 4) | 16.22 (19.05, 12.50) | 6 (67) | 3, 3 | 8.959 (6.361, 30.295) |
| *Brachyphylla cavernarum* | 64 (17, 47) | 24 (6, 18) | 37.5 (35.29, 39.13) | 7 (93) | 3, 4 | 7.492 (7.029, 15.332) |
| **Culebrones Cave Total** | **1,018 (519, 499)** | **306 (162, 144)** | **30.06 (31.21, 28.86)** | **37 (85)** | **9, 28** | **43.394 (38.369, 66.871)** |
| *Artibeus jamaicensis* | 57 (33, 24) | 24 (14, 10) | 42.11 (42.42, 41.67) | 7 (61) | 2, 5 | 11.421 (7.486, 47.238) |
| *Eptesicus fuscus* | 11 (0, 11) | 0 (NA, 0) | NA | NA | NA | NA |
| **Larva Cave Total** | **68 (33, 35)** | **24 (14, 10)** | **35.29 (42.42, 28.57)** | **7 (61)** | **2, 5** | **11.434 (7.487, 47.348)** |
| **Grand Total** | **1,086 (552, 534)** | **330 (176, 154)** | **30.39 (31.88, 28.84)** | **41 (80)** | **11, 30** | **51.116 (43.262, 86.239)** |

### OTU Delineation

Both alignment techniques resulted in two similar, clearly differentiated potential cutoff ranges, as demonstrated by consistent peaks in the histograms (Figure 1). The lower level of taxonomic grouping of OTUs was defined as those with a cutoff between 90 – 92%. This means that two sequences with less than 90% genetic similarity would be considered to be different OTUs. The higher level of taxonomic grouping was defined as those with a cutoff between 74 and 80%. As such, two sequences with less than 74% genetic similarity would be considered to be different OTUs. The number of OTUs resulting from the cutoff extremes for both levels of classification, when total alignment and pairwise alignment were considered (74, 81, 90 and 93 PID cutoffs; Figure 1), were 29, 31, 42, and 47 OTUs, respectively.

Affinity propagation resulted in 45 OTUs. Although OTU richness did not equal any values generated using the PID histograms, 45 OTUs is within the range of OTUs generated using the lower-level taxonomic cutoff (i.e. 42 – 47 OTUs at 90 – 93% PID cutoff), suggesting this cutoff is more strongly supported by affinity propagation than is the higher-level cutoff. Additionally, no two described herpesvirus species were greater than 90% similar. However, Ateline herpesvirus 3 and Saimirine herpesvirus 2, two gammaherpesvirus species, and Cercopithecine herpesvirus 5 and Macacine herpesvirus 3, two betaherpesvirus species, differed by 83.6% and 78.5%, respectively. Additionally, two strains of Human herpesvirus 6, a betaherpesvirus, and Human herpesvirus 4, a gammaherpesvirus, differed by 96.2 and 99.1%, respectively. These PIDs of established viral species further support the use of the lower-level taxonomic cutoff.

To refine the exact number of OTUs, two neighbor-joining trees were generated from the exemplar sequences, as determined using affinity propagation, one for the gammaherpesviruses and one for the betaherpesviruses. (Figure 2). When exemplar sequences were greater than 90% similar (the most conservative cutoff within the lower taxonomic range), clusters were merged to represent a single OTU. Additionally, each OTU represented by a single occurrence (i.e. a single clone sequence) was examined individually. If the singleton sequence was greater than 74% similar to the exemplar of another OTU, the least conservative cutoff point demonstrated by PID histograms, it was clustered with that OTU. Singleton sequences less than or equal to 74% similar to each of the other exemplars were considered to be unique OTUs. After refinement, 41 OTUs remained. Preliminary data collected from Mata de Plátano Nature Reserve in 2016 corroborated the existence of OTUs 12, 17, and 25. These OTUs were found in four host individuals in 2016, representing host species *Mor. blainvillii* (OTUs 12 and 25; two individuals) and *A. jamaicensis* (OTU 17; two individuals). All OTUs found in 2016 were detected in 2017.

**Figure 2:**
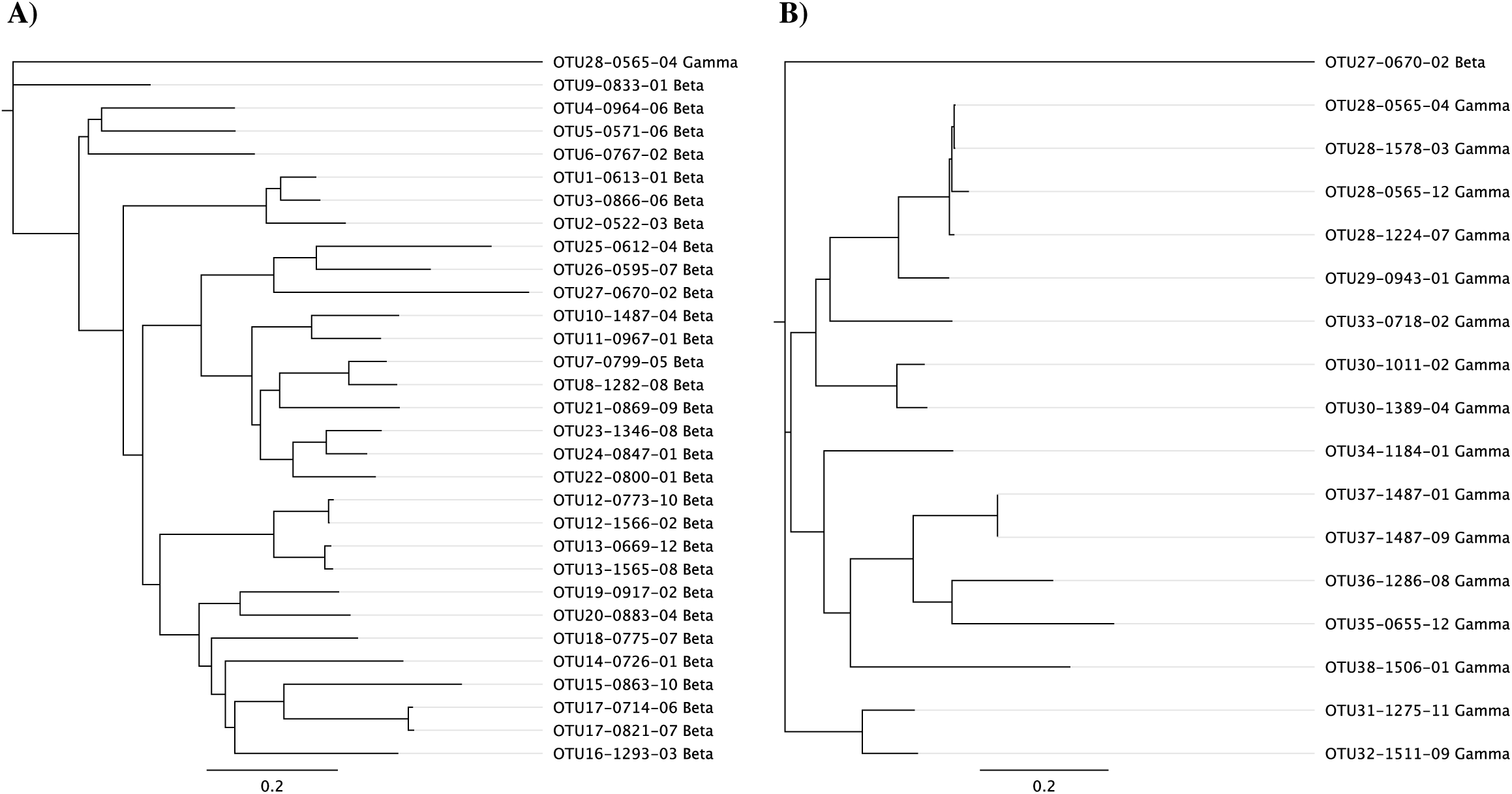
Neighbor-joining trees of betaherpesvirine (a) and gammaherpesvirine (b) exemplars, as defined using affinity propagation. OTUs are named to reflect manual refinement of OTUs after affinity propagation. For example, affinity propagation separated OTU 28, a gammaherpesvirus, into four separate clusters. However, the similarity of the exemplars suggested that these clusters should all be grouped into a single OTU. These trees provide biological support for the manual refinement of OTUs. OTUs 39-41, all betaherpesviruses, were not included in the tree. These OTUs were represented by three highly divergent sequences (< 50% genetic similarity), and their inclusion in the tree impaired visualization of all other relationships.

No OTU existed at a prevalence greater than 6.2% in the entire host community (i.e. across all host species). When viral metacommunities were examined at the level of host population (i.e. in a single species), however, OTU prevalence reached a peak of 26.3% (OTU 17 in *A. jamaicensis*; Table 2). At the population-level, four OTUs reached a prevalence greater than 20%, and 29 OTUs existed at prevalence values between 1% and 10%. Five OTUs existed at a prevalence less than 1% in all of their host populations. OTU richness per host population ranged from three to 15. In comparison, humans are the only host species for which comprehensive data are available on total herpesvirus richness, and only eight known herpesvirus species infect humans, despite their cosmopolitan distribution.

**Table 2:** Percent of hosts infected (prevalence) of each herpesvirus OTU at Mata de Plátano Nature Reserve (all bats), in each cave, and in each host population. Total number of sampled host individuals (N) is listed for each calculation. Continued on the next page.

| OTU | All Bats<br>(N = 1,086) | Culebrones<br>(N = 1,018) | Larva<br>(N = 68) | *Pteronotus quadridens*<br>(N = 288) | *P. portoricensis*<br>(N = 31) | *Mormoops blainvillii*<br>(N = 299) | *Monophyllus redmani*<br>(N = 262) | *Erophylla sezekorni*<br>(N = 74) | *Brachyphylla cavernarum*<br>(N = 64) | *Artibeus jamaicensis*<br>(N = 57) |
| --- | --- | --- | --- | --- | --- | --- | --- | --- | --- | --- |
| 1 | 2.762 | 2.947 | 0.000 | 10.07 | 3.226 | 0.000 | 0.000 | 0.000 | 0.000 | 0.000 |
| 2 | 1.565 | 1.670 | 0.000 | 5.208 | 0.000 | 0.334 | 0.382 | 0.000 | 0.000 | 0.000 |
| 3 | 0.829 | 0.884 | 0.000 | 3.125 | 0.000 | 0.000 | 0.000 | 0.000 | 0.000 | 0.000 |
| 4 | 1.197 | 1.277 | 0.000 | 3.472 | 0.000 | 0.669 | 0.382 | 0.000 | 0.000 | 0.000 |
| 5 | 0.552 | 0.589 | 0.000 | 2.083 | 0.000 | 0.000 | 0.000 | 0.000 | 0.000 | 0.000 |
| 6 | 0.276 | 0.295 | 0.000 | 1.042 | 0.000 | 0.000 | 0.000 | 0.000 | 0.000 | 0.000 |
| 7 | 0.276 | 0.196 | 1.429 | 0.000 | 0.000 | 0.000 | 0.763 | 0.000 | 0.000 | 1.754 |
| 8 | 0.184 | 0.196 | 0.000 | 0.000 | 0.000 | 0.000 | 0.763 | 0.000 | 0.000 | 0.000 |
| 9 | 0.092 | 0.000 | 1.429 | 0.000 | 0.000 | 0.000 | 0.000 | 0.000 | 0.000 | 1.754 |
| 10 | 0.276 | 0.295 | 0.000 | 0.000 | 0.000 | 0.000 | 0.000 | 4.054 | 0.000 | 0.000 |
| 11 | 0.092 | 0.098 | 0.000 | 0.000 | 0.000 | 0.000 | 0.000 | 1.351 | 0.000 | 0.000 |
| 12 | 5.801 | 5.992 | 2.857 | 0.694 | 0.000 | 19.73 | 0.000 | 0.000 | 0.000 | 3.509 |
| 13 | 3.683 | 3.929 | 0.000 | 0.347 | 0.000 | 13.04 | 0.000 | 0.000 | 0.000 | 0.000 |
| 14 | 0.368 | 0.000 | 5.714 | 0.000 | 0.000 | 0.000 | 0.000 | 0.000 | 0.000 | 7.012 |
| 15 | 0.276 | 0.294 | 0.000 | 0.000 | 0.000 | 0.000 | 0.763 | 0.000 | 1.563 | 0.000 |
| 16 | 0.092 | 0.098 | 0.000 | 0.000 | 0.000 | 0.000 | 0.382 | 0.000 | 0.000 | 0.000 |
| 17 | 1.473 | 0.098 | 21.43 | 0.000 | 0.000 | 0.000 | 0.382 | 0.000 | 0.000 | 26.31 |
| 18 | 0.092 | 0.098 | 0.000 | 0.000 | 0.000 | 0.000 | 0.000 | 1.351 | 0.000 | 0.000 |
| 19 | 0.460 | 0.491 | 0.000 | 0.000 | 0.000 | 0.000 | 0.000 | 0.000 | 7.813 | 0.000 |
| 20 | 0.645 | 0.688 | 0.000 | 0.000 | 0.000 | 0.000 | 0.000 | 0.000 | 10.93 | 0.000 |
| 21 | 1.289 | 1.375 | 0.000 | 0.000 | 0.000 | 0.000 | 5.343 | 0.000 | 0.000 | 0.000 |
| 22 | 0.276 | 0.295 | 0.000 | 0.000 | 0.000 | 0.000 | 0.000 | 4.054 | 0.000 | 0.000 |
| 23 | 0.645 | 0.688 | 0.000 | 0.000 | 0.000 | 0.000 | 0.000 | 0.000 | 10.94 | 0.000 |
| 24 | 0.276 | 0.295 | 0.000 | 0.000 | 0.000 | 0.000 | 0.000 | 0.000 | 4.688 | 0.000 |
| 25 | 2.855 | 3.045 | 0.000 | 0.694 | 0.000 | 9.699 | 0.000 | 0.000 | 0.000 | 0.000 |
| 26 | 0.552 | 0.589 | 0.000 | 0.347 | 0.000 | 1.672 | 0.000 | 0.000 | 0.000 | 0.000 |
| 27 | 0.552 | 0.589 | 0.000 | 0.000 | 0.000 | 2.007 | 0.000 | 0.000 | 0.000 | 0.000 |
| 28 | 6.169 | 6.582 | 0.000 | 0.347 | 0.000 | 0.334 | 24.81 | 0.000 | 0.000 | 0.000 |
| 29 | 0.276 | 0.295 | 0.000 | 0.000 | 0.000 | 0.000 | 1.145 | 0.000 | 0.000 | 0.000 |
| 30 | 1.289 | 1.375 | 0.000 | 0.347 | 0.000 | 0.000 | 0.000 | 0.000 | 20.31 | 0.000 |
| 31 | 0.184 | 0.196 | 0.000 | 0.000 | 6.452 | 0.000 | 0.000 | 0.000 | 0.000 | 0.000 |
| **32** | **0.092** | **0.098** | 0.000 | **0.347** | 0.000 | 0.000 | 0.000 | 0.000 | 0.000 | 0.000 |
| **33** | **1.197** | 0.000 | **18.57** | 0.000 | 0.000 | 0.000 | 0.000 | 0.000 | 0.000 | **22.81** |
| **34** | **0.737** | **0.786** | 0.000 | **0.347** | 0.000 | 0.000 | 0.000 | **9.459** | 0.000 | 0.000 |
| **35** | **0.184** | **0.196** | 0.000 | 0.000 | 0.000 | 0.000 | 0.000 | 0.000 | **3.125** | 0.000 |
| **36** | **0.552** | **0.589** | 0.000 | 0.000 | 0.000 | 0.000 | **2.290** | 0.000 | 0.000 | 0.000 |
| **37** | **0.092** | **0.098** | 0.000 | 0.000 | 0.000 | 0.000 | 0.000 | **1.351** | 0.000 | 0.000 |
| **38** | **0.184** | **0.196** | 0.000 | 0.000 | **6.451** | 0.000 | 0.000 | 0.000 | 0.000 | 0.000 |
| **39** | **0.092** | 0.000 | **1.429** | 0.000 | 0.000 | 0.000 | 0.000 | 0.000 | 0.000 | **1.754** |
| **40** | **0.092** | **0.098** | 0.000 | **0.347** | 0.000 | 0.000 | 0.000 | 0.000 | 0.000 | 0.000 |
| **41** | **0.092** | **0.098** | 0.000 | 0.000 | 0.000 | **0.334** | 0.000 | 0.000 | 0.000 | 0.000 |

### Viral Discovery Curves

The Chao2 estimate of herpesvirus richness in the bats of both caves was 51.1 OTUs (Table 1). As such, the detected OTUs (N = 41) represent about 80% of those in the entire herpesvirus metacommunity (Figure 3). Viral discovery curves were more saturated in Culebrones (85%) than in Larva (61%) Cave, and host species-level saturation ranged from 55% (*P. quadridens*) to 93% (*B. cavernarum*) (Table 1; Figure 3). A viral discovery curve was not generated for the herpesvirus metacommunity infecting *P. portoricensis*, as only three OTUs were detected.

**Figure 3:**
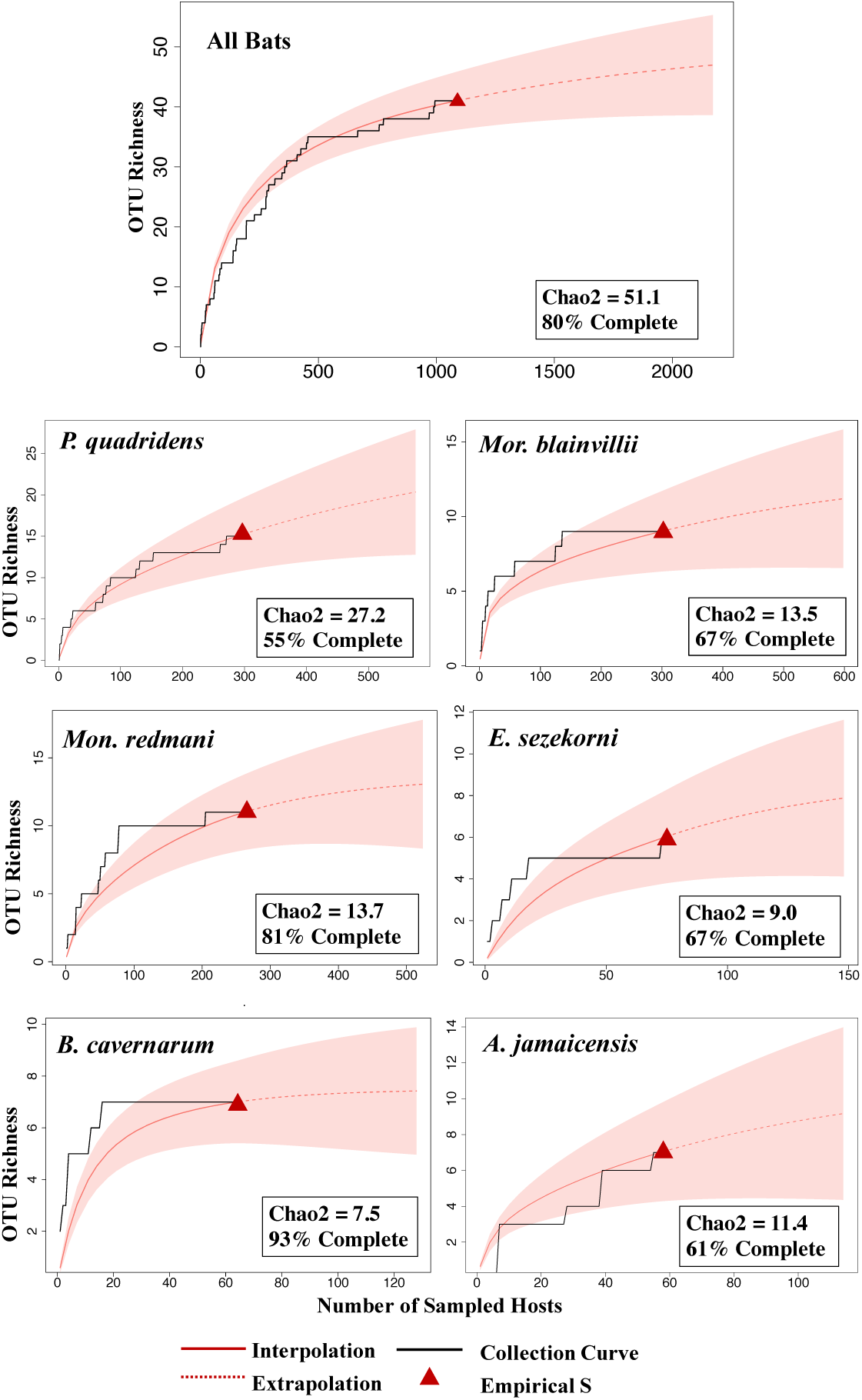
Viral discovery curves comparing number of detected herpesvirus (OTU richness) with number of sampled host bats for all bats at Mata de Plátano Nature Reserve (top panel) and for each host species (bottom six panels). The black lines represent the empirical pattern of OTU accumulation. The dashed red lines represent extrapolation of undiscovered herpesviruses in the metacommunities, with red shading demonstrating error associated with the estimates. Empirical viral richness per metacommunity (S) is depicted by red triangles. Chao2 numbers estimate true viral richness in each community and is used to calculate the percent of total estimated OTUs that were detected in samples (percent completeness). No discovery curve was generated for viruses infecting *P. portoricensis*, as only three OTUs were detected in this metacommunity.

### Host Preference

Thirteen OTUs were detected in one or more host species (Table 3). S_TD_* for each of these OTUs ranged from one (infecting two hosts from the same genus) to three (infecting three hosts of two families). Nine OTUs had significantly higher prevalence in a core host species than in all other host species combined (Table 4). Only two OTUs showed significantly higher prevalence in one host sex than in the other (Table 5). OTU 1 infected a significantly higher percentage of male than female *P. quadridens* (p = 0.021), and OTU 21 infected a significantly higher percentage of female than male *Mon. redmani* (p = 0.023).

**Table 3:** Host specificity index (S_TD_*) for each of the 13 herpesvirus OTUs that infect more than one host species. OTU prevalence is calculated for each host species, with number of sampled individuals in parentheses (N). Prevalence in the primary host species is bolded for each OTU.

| OTU | Number of Host Species Infected | Viral Sub-family | $S_{TD}^*$ | *Pteronotus quadridens* (N = 288) | *P. portoricensis* (N = 31) | *Mormoops blainvillii* (N = 299) | *Monophyllus redmani* (N = 262) | *Erophylla sezekorni* (N = 74) | *Brachyphylla cavernarum* (N = 64) | *Artibeus jamaicensis* (N = 57) |
| --- | --- | --- | --- | --- | --- | --- | --- | --- | --- | --- |
| 1 | 2 | Beta | 1.00 | **0.101** | 0.032 | 0.000 | 0.000 | 0.000 | 0.000 | 0.000 |
| 2 | 3 | Beta | 2.55 | **0.052** | 0.000 | 0.003 | 0.004 | 0.000 | 0.000 | 0.000 |
| 4 | 3 | Beta | 2.41 | **0.035** | 0.000 | 0.007 | 0.004 | 0.000 | 0.000 | 0.000 |
| 7 | 2 | Beta | 2.00 | 0.000 | 0.000 | 0.000 | 0.008 | 0.000 | 0.000 | **0.018** |
| 12 | 3 | Beta | 2.84 | 0.007 | 0.000 | **0.197** | 0.000 | 0.000 | 0.000 | 0.035 |
| 13 | 2 | Beta | 2.00 | 0.004 | 0.000 | **0.130** | 0.000 | 0.000 | 0.000 | 0.000 |
| 15 | 2 | Beta | 2.00 | 0.000 | 0.000 | 0.000 | 0.008 | 0.000 | **0.016** | 0.000 |
| 17 | 2 | Beta | 2.00 | 0.000 | 0.000 | 0.000 | 0.004 | 0.000 | 0.000 | **0.263** |
| 25 | 2 | Beta | 2.00 | 0.007 | 0.000 | **0.097** | 0.000 | 0.000 | 0.000 | 0.000 |
| 26 | 2 | Beta | 2.00 | 0.004 | 0.000 | **0.017** | 0.000 | 0.000 | 0.000 | 0.000 |
| 28 | 3 | Gamma | 2.99 | 0.004 | 0.000 | 0.003 | **0.248** | 0.000 | 0.000 | 0.000 |
| 30 | 2 | Gamma | 3.00 | 0.004 | 0.000 | 0.000 | 0.000 | 0.000 | **0.188** | 0.000 |
| 34 | 2 | Gamma | 3.00 | 0.004 | 0.000 | 0.000 | 0.000 | **0.095** | 0.000 | 0.000 |

**Table 4:** For each of the 13 herpesvirus OTUs that infect more than one host species, comparison of OTU prevalence between core host species and all other host species for which it is found, using Fisher’s exact test. Significance at an alpha of 0.01 are marked with a single asterisk, while those significant at an alpha of 0.001 are marked with two asterisks.

| OTU | Number of Infected Individuals (Core Host) | Number of Infected Individuals (Satellite Hosts) | P-value |
| --- | --- | --- | --- |
| 1 | 29 | 1 | 0.21 |
| 2 | 15 | 1 | < 0.001** |
| 4 | 10 | 3 | 0.002* |
| 7 | 2 | 1 | 0.92 |
| 12 | 59 | 4 | < 0.001** |
| 13 | 39 | 1 | < 0.001** |
| 15 | 2 | 1 | 0.90 |
| 17 | 15 | 1 | < 0.001** |
| 25 | 29 | 2 | < 0.001** |
| 26 | 5 | 1 | 0.12 |
| 28 | 65 | 2 | < 0.001** |
| 30 | 12 | 1 | < 0.001** |
| 34 | 7 | 1 | < 0.001** |

**Table 5:**
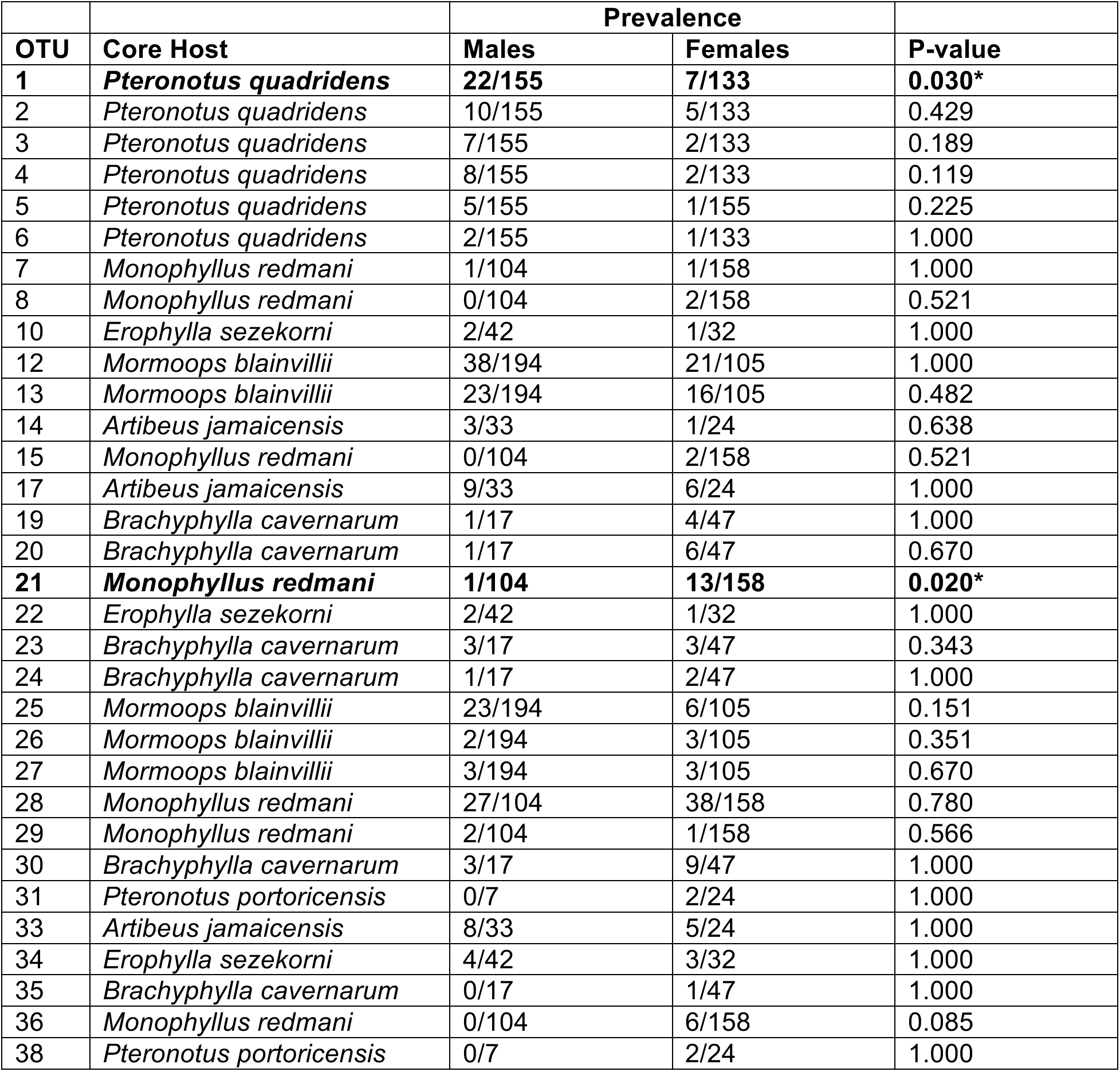
For the 32 herpesvirus OTUs infecting more than one host individual, prevalence is compared between males and females using Fisher’s exact test. The number of host individuals that tested positive is in the numerator, while the total number of captured host individuals is the denominator. Significance at the alpha level of 0.05 is shown with an asterisk, and rows with significant differences are bolded. For all OTUs that are not represented in this table, only a single individual tested positive.

## Discussion

Our extensive quantification of viral infection in bats represents the first comprehensive study of the virome of a wildlife host community and suggests that Puerto Rican bats offer a previously untapped resource for advancing the field of viral ecology. Even in the absence of full genome sequences, viral OTUs can be robustly differentiated and used to understand the natural history and host specificity of viruses.

The application of tools and perspectives from community ecology to better understand viral infection dynamics necessitates an objective, quantitatively-supported delineation of viral “species”. Officially designating novel viral species requires substantial data, oftentimes full genome sequences, and approval by the International Committee for the Taxonomy of Viruses (King et al. 2012). Consequently, OTUs are commonly used as proxies for species. Although multiple methods exist for delineating OTUs (e.g. histograms using percent genetic identity, Maes et al. 2009; or monophyletic groups, Anthony et al. 2015), subjectivity is generally required to choose the best cutoff point in terms of genetic similarity (i.e. the histogram method) or evolutionary time (i.e. the monophyletic groups method). Affinity propagation is a machine-learning algorithm developed to cluster a broad range of data (e.g. images of faces, genes in a microarray, heavily travelled cities) and identify representative examples of each cluster (Frey and Dueck 2007). The method has been used to cluster strains of rabies viruses (Fischer et al. 2017), but has never been used to identify viral OTUs for analyses in the context of community ecology. Here, we show that affinity propagation can be used as a quantitative complement, in congruence with currently defined herpesvirus species, to objectively support current methods of viral OTU delineation.

A limit to working with viruses in an ecological context is the short viral lifespan or activation cycle (e.g. lytic stage), as compared to that in free-living or macroparasitic organisms. Short lifespans affect temporal turnover of detected OTUs and could influence interpretation of results. Although it is common for some viral infections to last only hours or days, the lytic cycle of herpesvirus infections is typically longer than that of most other viruses (King et al. 2012). This represents an advantage in the use of herpesviruses for preliminary exploration of viral ecology, as the duration during which OTUs can be detected will be greater than that of other taxa. Indeed, all OTUs detected during preliminary sampling in 2016 (i.e. OTUs 12, 17, and 25) were detected again in 2017, in the same core host species (Table 2).

Another advantage to studying herpesviruses from an ecological context is that this viral family often exists at high prevalence levels (Cone et al. 1993, Kidd et al. 1996, Cortez et al. 2008, Imbronito et al. 2008, Tenorio de Franca et al. 2012, Phalen et al. 2017, Tazikeh et al. 2019). While some surveys (e.g. Cone et al. 1993, Kidd et al. 1996, Phalen et al. 2017) show prevalence values greater than 70% for individual viral species, the population-level prevalence levels of OTUs in Puerto Rican bats are consistent with other surveyed populations of healthy humans (Tenorio de Franca et al. 2012) and non-human animals (Corez et al. 2008, Tazikeh et al. 2019).

Generally, herpesviruses are highly host specific, but when host-virus relationships are examined at an evolutionary timescale, host switching has been the norm (Escalera-Zamudio et al. 2016, Azab et al. 2018). As such, we expected herpesviruses that infected multiple host species to have core hosts. Overall, we found evidence of host sharing among bat species at Mata de Plátano Nature Reserve, regardless of cave. The existence of host sharing is further supported by viral prevalence and viral discovery curves. A higher prevalence at the population-level (i.e. a single species) than the community-level, as seen here (Table 2), is consistent with host-specificity. Also, at the largest scale (i.e. the entire nature reserve), sampling completeness reached 80%. This is in contrast to accumulation curves for viral metacommunities infecting particular host populations, which were as low as 55% (*P. quadridens*). This suggests that undiscovered OTUs at the scale of host population have already been discovered at the scale of nature reserve (i.e. have already been discovered in another host species). This supports the hypothesis that certain OTUs that are common in a core host are rare in all other host species. Indeed, nine of the 13 multi-host OTUs had significantly higher prevalence in a single host than in all other hosts in which it occurred combined (Table 3). Those that did not have significantly different prevalence were rare OTUs in all infected hosts (OTU 7, OTU 15, OTU 26), or were compared between host species with drastically different sample sizes (OTU 1, found in *P. quadridens* and *P. portoricensis*; with 288 and 31 sampled individuals, respectively).

In conclusion, ecological tools and perspectives can be used to understand viral infection and transmission, generating a multitude of hypotheses in the emerging field of viral ecology. Future studies with goals of monitoring public health should increase the number of sampled individuals for each host species to enhance the likelihood of uncovering patterns of host preference and reservoir status. Studies aimed at viral ecology will likely require focus on more prevalent viral taxa, such as herpesviruses, instead of rare viral taxa of more zoonotic concern, such as coronaviruses or filoviruses. Further ecological research should prioritize sampling host individuals from multiple field locations, addressing viral infection and transmission at multiple spatial scales, while decoupling the spatial and ecological mechanisms that shape viral community assembly.

## Acknowledgments

We gratefully acknowledge Armando Rodriguez-Duran for invaluable logistical help in the field, as well as the intrepid field team, including Sarah Stankavich, Jose Rivera, Melanie Hodge, Brian Springall, Elspeth Pierce, Emily Stanford, Juan Contreras, Ismael Ribot, and Frank Sjodin. Isamara Navarrete-Macias and Eliza Liang trained ARS on laboratory methods, and Heather Wells provided invaluable lab and bioinformatics help. Cale Basaraba provided coding assistance. We also thank Janine Caira, Sarah Knutie, and Mark Urban for constructive discussions that advanced the intellectual development of the work. ARS was funded by the National Science Foundation Graduate Research Fellowship Program and a Jorgensen Fellowship from the University of Connecticut. MRW was funded by a grant from NSF (DEB-1546686 and DEB-1831952). Fieldwork was supported by Sigma Xi; the Royal Society of Tropical Medicine and Hygiene; and the University of Connecticut’s Department of Ecology and Evolutionary Biology, El Instituto via the Tinker Foundation, and the Office of the Vice President for Research. This project also benefited from support from the USAID PREDICT project.

## References

Anthony SJ, Epstein JH, Murray KA, Navarrete-Macias I, Zambrana-Torrelio CM, Solovyov A, Ojeda-Flores R, Arrigo NC, Islam A, Khan SA, Hosseini P, Bogich TL, Olival KJ, Sanchez-Leon MD, Karesh WB, Goldstein T, Luby SP, Morse SS, Mazet JAK, Daszak P, Lipkin WI (2013) A strategy to estimate unknown viral diversity in mammals. mBio 4: e00598–13.

Anthony SJ, Islam A, Johnson C, Navarrete-Macias I, Liang E, Jain K, Hitchens PL, Che X, Soloyvov A, Hicks AL, Ojeda-Flores R, Zambrana-Torrelio C, Ulrich W, Rostal MK, Petrosov A, Garcia J, Haider N, Wolfe N, Goldstein T, Morse SS, Rahman M, Epstein JH, Mazet JK, Daszak P, Lipkin WI (2015) Non-random patterns in viral diversity. Nature Communications 6: 8147.

Anthony SJ, Johnson CK, Greig DJ, Kramer S, Che X, Wells H, Hicks AL, Joly DO, Wolfe ND, Daszak P, Karesh W, Lipkin WI, Morse SS, PREDICT Consortium, Mazet JAK, Goldstein T (2017) Global patterns in coronavirus diversity. Virus Evolution 3: vex012.

Azab W, Dayaram A, Greenwood AD, Osterrieder N (2018) How host specific are herpesviruses? Lessons from herpesviruses infecting wild and endangered mammals. Annual Review of Virology 5: 53–68.

Bodenhofer U, Kothmeier A, Hochreiter S (2011) APCluster: a R package for affinity propagation clustering. Bioinformatics 17: 2463–2464.

Bush AO, Holmes JC (1986) Intestinal helminths of lesser scaup ducks: an interactive community. Canadian Journal of Zoology 64: 142–152.

Chao A (1987) Estimating the population size for capture-recapture data with unequal catchability. Biometrics 43: 783–791.

Cortez PP, Carvalheira J, Pauperio S, Thompson G (2008) Prevalence of ovine herpesvirus type 2 in north-west Portugal. Veterinary Record 162: 282–284.

De Benedictis P, Schultz-Cherry S, Burnham A, Cattoli G (2011) Astrovirus infections in humans and animals – molecular biology, genetic diversity, and interspecies transmissions. Infection, Genetics and Evolution 11: 1529–1544.

Escalera-Zamudio M, Rojas-Anaya E, Kolokotronis SO, Taboada B, Loza-Rubio E, Mendez-Ojeda ML, Arias CF, Osterrieder N, Greenwood AD (2016) Bats, primates, and the evolutionary origins and diversification of mammalian Gammaherpesviruses. mBio 7: e01425–16.

Fischer S, Freuling CM, Muller T, Pfaff F, Bodenhofer U, Hoper D, Fischer M, Marston DA, Fooks AR, Mettenleiter TC, Conraths FJ, and Homeier-Bachmann T (2017) Defining objective clusters for rabies virus sequences using affinity propagation clustering. PLoS Neglected Tropical Diseases 12: e0006182.

Frey BJ and Dueck D (2007) Clustering by passing messages between data points. Science 315: 972–976.

Gannon MR, Kurta A, Rodríguez-Duran A, and Willig MR (2005) Bats of Puerto Rico: An Island Focus and a Caribbean Perspective. Lubbock, TX: Texas Tech University Press.

Greenblatt RJ, Quackenbush SL, Casey RN, Rovnak J, Balazs GH, Work TM, Casey JW, Sutton CA (2005) Genomic variation of the fibropapilloma-associated marine turtle herpesvirus across seven geographic areas and three host species. Journal of Virology 79: 1125–1132.

Henaux V, Samuel MD (2011) Avian influenza shedding patterns in waterfowl: implications for surveillance, environmental transmission, and diseases spread. Journal of Wildlife Diseases 47: 566–578.

Holz PH, Lumsden LF, Druce J, Legione AR, Vaz P, Devlin JM, Hufschmid J (2018) Virus survey in populations of two subspecies of bent-winged bats (*Miniopterus orianae bassanii* and *oceanensis*) in south-eastern Australia reveals a high prevalence of diverse herpesviruses. PLoS ONE 13: e0197625.

Hsieh TC, Ma KH, Chao A (2016) iNEXT: an R package for rarefaction and extrapolation of species diversity (Hill numbers). Methods in Ecology and Evolution 7: 1451–1456.

Jolles AE, Ezenwa VO, Etienne RS, Turner WC, Olff H (2008) Interactions between macroparasites and microparasites drive infection patterns in free-ranging African buffalo. Ecology 89: 2239–2250.

King AMQ, Adams MJ, Carstens EB, Lefkowitz EJ (2012) Virus Taxonomy: Classification and Nomenclature of Viruses. Ninth report of the International Committee on Taxonomy of Viruses, San Diego, CA: Academic Press.

Klenk K, Snow J, Morgan K, Bowen R, Stephens M, Foster F, Gordy P, Beckett S, Komar N, Gubler D, Bunning M (2004) Alligators as West Nile virus amplifiers. Emerging Infectious Diseases 10: 2150–2155.

Lee N, Chan PKS, Hui DSC, Rainer TH, Wong E, Choi KW, Lui GCY, Wong BCK, Wong RYK, Lam WY, Chu IMT, Lai RWM, Cockram CS, Sung JJY (2009) Viral loads and duration of viral shedding in adult patients hospitalized with influenza. The Journal of Infectious Diseases 200: 492–500.

Maes P, Klempa B, Clement J, Matthijnssens J, Gajdusek DC, Kruger DH, Van Ranst M (2009) A proposal for new criteria for the classification of hantaviruses, based on S and M segment protein sequences. Infection, Genetics and Evolution 9: 813–820.

Pages H, Aboyoun P, Gentleman R, DebRoy S (2017) Biostrings: Efficient manipulation of biological strings. R package version 2.46.0.

Pedersen AB, Fenton A (2006) Emphasizing the ecology in parasite community ecology. Trends in Ecology and Evolution 22: 133–139.

Phalen DN, Alvarado C, Grillo V, Mason P, Dobson E, Holz P (2017) Prevalence of columbid herpesvirus infection in feral pigeons from New South Wales and Victoria, Australia, with spillover into a wild powerful owl (*Ninox struena*). Journal of Wildlife Diseases 53: 543–551.

Poulin R, Mouillot D (2005) Combining phylogenetic and ecological information into a new index of host specificity. Journal of Parasitology 91: 511–514.

Pozo F, Juste J, Vazquez-Moron S, Aznar-Lopez C, Ibanez C, Garin I, Aihartza J, Casas I, Tenorio A, Echevarria JE (2016) Identification of novel betaherpesviruses in Iberian bats reveals parallel evolution. PLoS ONE 11: e0169153.

Razafindratsimandresy R, Jeanmaire EM, Counor D, Vasconcelos PF, Sall AA, Reynes JM (2009) Partial molecular characterization of alphaherpesviruses isolated from tropical bats. Journal of General Virology 90: 44–47.

Rodríguez-Duran, A (2009) Bat assemblages in the West Indies: the role of caves. In Island Bats: Evolution, Ecology, and Conservation, Fleming TH and Racey PA (editors) Chicago, IL: University of Chicago Press, pp 265–280.

Russell GC, Stewart JP, Haig DM (2009) Malignant catarrhal fever: a review. The Veterinary Journal 179: 324–335.

Sasaki M, Setiyono A, Handharyani E, Kobayashi S, Rahmadani I, Taha S, Adiani S, Subangkit M, Nakamura I, Sawa H, Kimura T (2014) Isolation and characterization of a novel alphaherpesvirus in fruit bats. Journal of Virology 88: 9819–9829.

Seabloom EW, Borer ET, Gross K, Kendig AE, Lacroix C, Mitchell CE, Modrecai EA, Power AG (2015) The community ecology of pathogens: coinfection, coexistence and community composition. Ecology Letters 18: 401–415.

Tandler B (1996) Cytomegalovirus in the principal submandibular gland of the little brown bat, *Myotis lucifugus*. Journal of Comparative Pathology 114: 1–9.

Tang XC, Zhang JX, Zhang SY, Wang P, Fan XH, Li LF, Li G, Dong BQ, Liu W, Cheung CL, Xu KM, Song WJ, Vijaykrishna D, Poon LLM, Peiris JSM, Smith GJD, Chen H, Guan Y (2006) Prevalence and genetic diversity of coronaviruses in bats from China. Journal of Virology 80: 7481–7490.

Tazikeh A, Raoofi A, Madadgar O, Akbarein H, Mashhadi AG. 2019. A survey of equine herpes virus 4 infection in four provinces of Iran using Real Time PCR Taqman assay. Journal of Veterinary Research 73: 483–490.

Telfer S, Lambin X, Birtles R, Beldomenico P, Burthe S, Paterson S, Begon M (2010) Species interactions in a parasite community drive infection risk in a wildlife population. Science 330: 243–246.

Van De Vanter DR, Warrener P, Bennett L, Schultz ER, Coulter S, Garber RL, Rose TM (1996) Detection and analysis of diverse herpesviral species by consensus primer PCR. Journal of Clinical Microbiology 34: 1666–1671.

Vetter P, Fischer II WA, Schibler M, Jacobs M, Bausch DG, Kaiser L (2016) Ebola virus shedding and transmission: review of current evidence. The Journal of Infectious Diseases 214: S177–S184.

Wacharapluesadee S, Boongrid K, Wanghongsa S, Ratanasetyuth N, Supavongwong P, Saengsen D, Gongal GN, Hemachudha T (2010) A longitudinal study of the prevalence of Nipah virus in *Pteropus lylei* bats in Thailand: evidence for seasonal preference in disease transmission. Vector-Borne and Zoonotic Diseases 10: 183–190.

Wada Y, Sasaki M, Setiyono A, Handharyani E, Rahmadani I, Taha S, Adiani S, Latief M, Kholilullah ZA, Subangkit M, Kobayashi S, Nakamura I, Kimura T, Orba Y, Sawa H (2018) Detection of novel gammaherpesviruses from fruit bats in Indonesia. Journal of Medical Microbiology 67: 415–422.

Wassertheil-Smoller S, Smoller J (2015) Biostatistics and Epidemiology: A Primer for Health and Biomedical Professionals Fourth Edition, New York, NY: Springer.

Weigler BJ (1992) Biology of B virus in macaque and human hosts: a review. Clinical Infectious Diseases 14: 555–567.

Wibbelt G, Kurth A, Yasmum N, Bannert M, Nagel S, Nitsche A, Ehlers B (2007) Discovery of herpesviruses in bats. Journal of General Virology 88: 2651–2655.

Wibbelt G, Moore MS, Schountz T, Voigt CC (2010) Emerging diseases in Chiroptera: why bats? Biology Letters 6: 438–440.

Winker K, Spackman E, Swayne DE. Rarity of Influenza A virus in spring shorebirds, southern Alaska. Emerging Infectious Diseases 14: 1314–1316.

Zhang H, Todd S, Tachedijian M, Barr JA, Luo M, Yu M, Marsh GA, Crameri G, Wang L (2012) A novel bat herpesvirus encodes homologues of major histocompatibility complex class I and II, c-type lectin, and a unique family of immune-related genes. Journal of Virology 86: 8014–8030.

Zheng X, Qiu M, Chen S, Ziao J, Ma L, Liu S, Zhou J, Zhang Q, Li X, Chen Z, Wu Y, Chen H, Jiang L, Xiong Y, Ma S, Zhong X, Huo S, Ge J, Cen S, Chen Q (2016) High prevalence and diversity of viruses in the subfamily Gammaherpesvirinae, family Herpesviridae, in fecal specimens from bats of different species in southern China. Archives of Virology 161: 135–140.

